# TREM2 deficiency causes region-specific brain effects in a mouse model of cerebral amyloid angiopathy

**DOI:** 10.64898/2026.04.17.719285

**Authors:** Constanza Mercado, Armando Amaro, Jonathan Martinez-Pinto, Ruben Vidal, Nur Jury-Garfe, Cristian A. Lasagna-Reeves

## Abstract

Cerebral amyloid angiopathy (CAA), a major vascular contributor to cognitive decline, is present in 85–95% of Alzheimer’s disease (AD) patients. Despite its high prevalence, the mechanisms by which CAA contributes to neurodegeneration remain poorly understood. Triggering receptor expressed on myeloid cells 2 (TREM2), an innate immune receptor expressed exclusively by microglia, regulates activation, phagocytosis, and amyloid clearance, thereby shaping neuroinflammation. Loss-of-function mutations in TREM2 markedly increase AD risk, but its role in CAA pathology remains unknown. To investigate this, we crossed the Familial Danish Dementia (Tg-FDD) mouse model, which accumulates robust vascular amyloid, with TREM2 knockout (TREM2KO) mice to generate Tg-FDD/TREM2KO animals. Histological and transcriptomic analyses revealed region-specific effects of TREM2 deficiency. In the cortex, TREM2 loss markedly reduced vascular amyloid deposition, accompanied by decreased tau pathology. In contrast, in the cerebellum, TREM2 deletion exacerbated vascular amyloid accumulation, promoted astrogliosis, and enhanced tau pathology. Transcriptomic profiling further identified distinct neuroinflammatory signatures between cortex and cerebellum, particularly in cytokine signaling, matrix remodeling, and lipid metabolism. Together, these findings demonstrate that TREM2 deficiency leads to region-specific effects on CAA, revealing extensive regional variability in vascular amyloid pathology and underscoring the importance of considering these differences when developing TREM2-based therapies.

## INTRODUCTION

Cerebral amyloid angiopathy (CAA) is characterized by the deposition of amyloid in cerebral blood vessels. Although closely related to Alzheimer’s disease (AD) at the molecular level, it remains clinically distinct (1). Vascular amyloid accumulation is detected in approximately 85–95% of individuals with AD (1,2), highlighting CAA as a major vascular contributor to age-related cognitive decline. The mechanisms underlying CAA pathogenesis and its downstream effects on the brain are complex and remain incompletely understood. While CAA is primarily associated with beta-amyloid (Aβ) accumulation (2), other amyloid proteins can also deposit in cerebral blood vessels (3). This suggests that CAA represents a group of biochemically and genetically diverse disorders, unified by the presence of amyloid deposits in the walls of cerebral arteries and, in some cases, in the capillaries of the central nervous system (CNS) parenchyma and leptomeninges.

Interestingly, variants in Triggering Receptor Expressed on Myeloid Cells 2 (TREM2), a microglial receptor in the brain, have been identified as increasing the risk of developing late-onset AD (LOAD) (4,5). Functionally, TREM2 regulates several key microglial processes, including phagocytosis, chemotaxis, and cell survival. These functions are essential for maintaining brain homeostasis and contribute to protection against a wide range of CNS pathologies, including cerebrovascular diseases and neurodegenerative disorders (6). Numerous studies have explored how TREM2 influences amyloid and tau aggregation across different AD mouse models, revealing that TREM2 deficiency can exert either protective or detrimental effects depending on the context (7–16). Despite these advances, the specific role of TREM2 in CAA remains poorly understood.

We previously showed that the transgenic mouse model of familial Danish dementia (Tg-FDD), which develops exclusively CAA and exhibits vascular deposition of Danish amyloid (ADan) in vessels and leptomeninges (17), displays dysregulated TREM2 expression (18). This suggests that disrupted TREM2 homeostasis may play a key role in CAA pathogenesis.

Therefore, this study aims to determine whether TREM2 deficiency exerts an effect on the amyloid-associated vascular environment in the Tg-FDD mouse model. Our finding indicates that loss of TREM2 in the Tg-FDD mouse led to distinct regional outcomes. In the cortex, we observed markedly reduced vascular amyloid deposition, accompanied by decreased tau oligomers. Conversely, in the cerebellum, TREM2 deletion exacerbated vascular amyloid accumulation and was associated with increased astrogliosis and enhanced tau oligomers. Transcriptomic profiling further revealed divergent neuroinflammatory signatures between the cortex and cerebellum, particularly in pathways related to cytokine signaling, extracellular matrix remodeling, and lipid metabolism. Together, these findings demonstrate that TREM2 deficiency produces region-specific effects on CAA, highlighting substantial regional variability in vascular amyloid pathology and emphasizing the importance of accounting for these differences in the development of TREM2-targeted therapies.

## MATERIALS AND METHODS

### Transgenic mouse model

Experiments were conducted using C57BL/6J (WT) (JAX stock #000664), Tg-FDD, Tg-FDD/TREM2KO, and TREM2KO (JAX stock #027197) mice. All mice were housed at the Indiana University School of Medicine (IUSM) animal care facility and maintained under USDA standards (12-h light/dark cycle, food and water ad libitum), in accordance with the Guide for the Care and Use of Laboratory Animals (National Institutes of Health, Bethesda, MD). Animals were anesthetized and euthanized following protocols approved by the IUSM Institutional Animal Care and Use Committee. Mice were deeply anesthetized prior to decapitation, and brains were collected and either stored at −80 °C or fixed in formalin as previously described (19,20). All experiments were performed using 22-month-old mice if otherwise mentioned.

### Brain sections immunofluorescence and Thioflavin-S staining

Paraffin-embedded sections were deparaffinized in xylene, rehydrated in ethanol (EtOH), and washed with deionized water. Antigen retrieval was performed by heating the sections in a high-pH solution for 12 min using a pressure cooker. After two washes in PBS (5 min each), sections were blocked for 1 h at room temperature (RT) in Animal-free blocker (SP-5035-100, Vector Laboratories) containing 0.3% Triton X-100. Primary antibodies anti-GFAP (Sigma, G3893), anti-IBA1 (Fujfilm, 019-19741), anti-αSMA (EMD Millipore, CBL171-l), anti-TOMA1 (EMD Millipore, MABN819) and anti-ADan 1699 (gift from Dr. Ruben Vidal) were applied in blocking solution and incubated overnight at 4 °C. The following day, sections were washed three times in PBS and incubated with secondary antibodies (1:500) for 2 h at RT. Vascular amyloid deposits were stained with 0.5% Thioflavin-S (Thio-S) for 15 min at RT, followed by four washes in PBS. Autofluorescence was quenched using TrueBlack (Biotium, 23007) for 3 min, after which sections were counterstained with TO-PRO™ for 30 min. Finally, sections were rinsed in PBS and mounted with Fluoromount aqueous mounting media (Sigma-Aldrich, F4680).

### Microscopy and image analysis

Pictures from the cortex and cerebellum were acquired using a 40× objective on a Leica DMi8 epifluorescence microscope equipped with LAS X software. Post-processing and fluorescence-intensity quantification were performed using ImageJ (National Institutes of Health, v1.53c). For thioflavin-S quantification, a region of interest (ROI) was drawn around the vasculature, delimited by αSMA, and applied to the thioflavin-S channel to calculate the percentage of immunoreactivity within the defined area (21). IBA1 and GFAP fluorescence intensities were quantified individually using ROIs placed around the vasculature within a 120 µm radius, measuring the Mean Gray Value (MGV) for each ROI (22). For TOMA1 staining, oligomers were quantified using the analyze particles function in ImageJ.

### RNA sequencing

Data used in this analysis, including library preparation, sequencing, sequence alignment, and gene count quantification, were obtained from our previous study (18). Nine-month-old Tg-FDD mice were compared with age-matched WT control. Differential expression (DE) was determined using a cutoff of fold change > 1.2 and an adjusted p-value < 0.05. These data were subsequently analyzed for genomic pathway analysis. Significant genes associated with the TREM2 pathway were visualized using RStudio (version 2025.05.01), and functional interactions and pathway enrichment analyses were performed using the STRING database to evaluate protein–protein interactions among TREM2-associated genes.

### NanoString gene expression analysis

Total messenger RNA (mRNA) was purified from the cortex and cerebellum of WT, Tg-FDD, and Tg-FDD/TREM2KO, mice and analyzed using the nCounter analysis system (NanoString Technologies, Seattle, WA, USA). The nCounter Neuroinflammation Profiling Panel, which included 770 genes covering core pathways of glial cell homeostasis and activation, was performed as previously described (21,23). Briefly, 100 ng of total RNA per sample (n = 3 per condition) were hybridized with probes for 16 h at 65 °C according to the manufacturer’s protocol. RNA quality was assessed using DV200 (%) and RNA Integrity Number (RIN) metrics (Supplementary table 1), and all samples analyzed had RIN values between 8.2 and 8.9. Counts for target genes were normalized to the most stable housekeeping genes as determined by nSolver software (v4.0) to account for variation in RNA content. The background signal was calculated as the mean value of the negative hybridization control probes. Expression data were excluded when signals were lower than the average of the negative controls, and probes with fewer than 100 reads in six or more samples were removed from the analysis. Downstream analyses and visualization of gene expression datasets were performed using nSolver software (v4.0), GraphPad Prism, and RStudio (version 2025.05.01).

### Reproducibility and Statistical analyses

Sample sizes were determined based on previous publications (21). The experimental analyses and data collection protocols were performed blind. Statistical analyses and graph generation were performed using GraphPad Prism and RStudio (version 2025.05.01). All data were first assessed for normality and outliers, followed by the appropriate statistical tests as indicated in the figure legends. Sex was considered as a biological variable in staining experiments. Data are presented as median ± standard error of the mean (SEM), unless otherwise stated. Statistical significance is indicated as follows: *p*<0.05 (*), *p*<0.01 (**), *p*<0.001 (***), *p*<0.0001 (****).

## RESULTS

### TREM2 pathway shows distinct regulation in the cortex and cerebellum of Tg-FDD mice

Previously, gene expression analysis revealed that several AD risk factors involved in immune response and lipid processing may also play a significant role in CAA, with dysregulation of TREM2 signaling potentially contributing to disease pathogenesis (18). Based on its established role in microglial function and AD progression (5,24), TREM2 emerged as a key candidate to investigate mechanisms underlying vascular amyloid pathology.

Upon ligand binding, TREM2 signals via Tyrobp, recruiting downstream molecules such as PLCγ2 and INPP5D, thereby initiating pathways that promote cell survival, proliferation, chemotaxis, and phagocytosis (25). To assess whether TREM2 signaling is altered in the Tg-FDD CAA model, we analyzed RNA-seq data from the cortex and cerebellum of 9-month-old Tg-FDD and WT mice, two brain regions characterized by the accumulation of vascular amyloid (17). Cerebellar data was previously reported (18), whereas cortical data is analyzed here for the first time, focusing specifically on the 25 genes comprising the TREM2 signaling pathway (26,27). Differential expression analysis revealed region-specific differences. In the cortex, one gene (*Cst7*) was significantly upregulated, while two genes (*Tyrob* and *Spp1*) were significantly downregulated in Tg-FDD compared to WT (Fig. 1A). In contrast, the cerebellum exhibited a more pronounced transcriptional response, with two genes (*Trem2,* and *Tyrobp*) that showed significant increases in expression and three genes (*Src*, *Ptk2b* and *Vav3*) that showed significant decreases in expression (Fig. 1B). To further illustrate these regional differences, a heatmap of the 25 TREM2 pathway genes was generated, highlighting distinct expression profiles between the cortex and cerebellum and confirming region-specific transcriptional regulation in Tg-FDD mice compared to WT (Fig. 1C and D).

**Figure 1:**
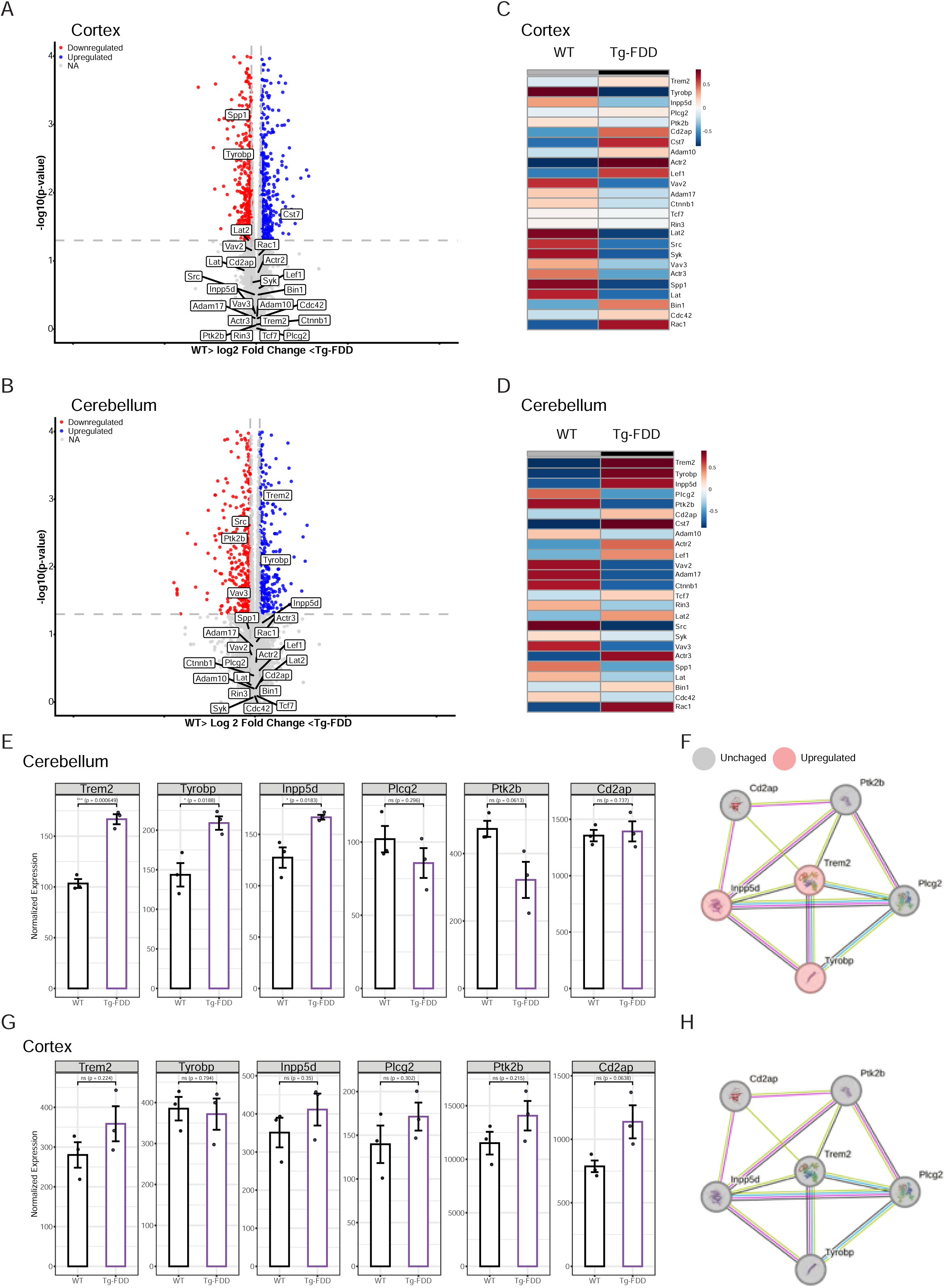
TREM2 pathway show distinct regulation in the cortex and cerebellum of Tg-FDD mice. Data reanalyzed from ref 17. A, B) Volcano plots showing differentially expressed genes of the TREM2 pathway. C, D) Heatmaps illustrating gene expression patterns (red = upregulated, blue = downregulated, NA = not assigned due to non-significant values). E, G) Normalized expression of TREM2 pathway–associated genes; F, H) corresponding network analysis of genes identified by RNA-Seq. Data are shown as mean ± SEM. Unpaired Student’s t-test, n = 3.

Then, we analyzed the expression of six selected AD risk genes related to the TREM2 pathway in both regions (26). In the cerebellum, three genes (*Trem2*, *Tyrobp*, and *Inpp5d*) were significantly upregulated, two genes (*Plcg2* and *Ptk2b*) showed a trend toward decreased expression, and *Cd2ap* showed no significant changes between Tg-FDD and WT mice (Fig. 1E and F). In contrast, in the cortex, no statistical differences were observed in any of the analyzed genes (Fig. 1G and H). Overall, these results indicate that TREM2 pathway genes are differentially regulated in Tg-FDD mice across brain regions, suggesting region-specific differences in the neuroinflammatory response associated with vascular amyloid pathology.

### TREM2 deficiency differentially modulates vascular amyloid in cortex and cerebellum of Tg-FDD mice

To evaluate whether reducing TREM2 levels produces region-specific effects in CAA, we analyzed vascular deposition in the Tg-FDD and Tg-FDD/TREM2KO mice using Thio-S staining in the cortex and cerebellum (Fig. 2A). In the cortex, Tg-FDD/TREM2KO mice exhibited a significant reduction in vascular Thio-S signal compared to Tg-FDD mice. In contrast, in the cerebellum, vascular amyloid burden was increased in Tg-FDD/TREM2KO mice vs Tg-FDD mice (Fig. 2B). These findings indicate that TREM2 deficiency has opposing effects on vascular amyloid deposition in the cortex and cerebellum in males as well as in female mice.

**Figure 2:**
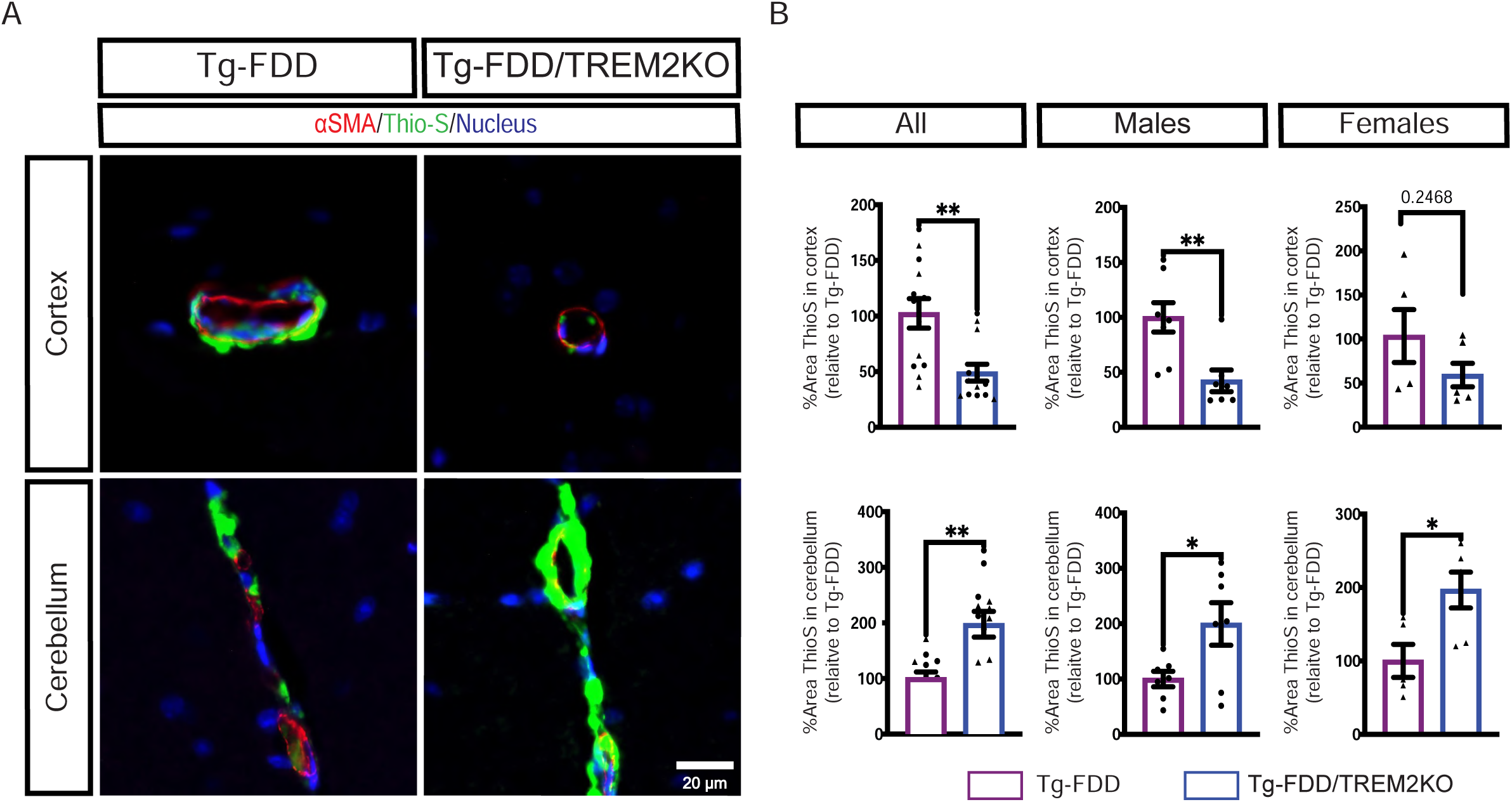
TREM2 deficiency differentially modulates vascular amyloid in cortex and cerebellum. A) Immunofluorescence for alpha smooth muscle actin (*α*SMA-red) and thioflavin-S staining (green) showing vascular amyloid deposition in the brain cortex and cerebellum of Tg-FDD and Tg-FDD/TREM2KO mice. Scale bar: 20 μm. B) Quantification of the percentage of thioflavin-S-positive surrounding area in the perivasculature. Data are shown as mean ± SEM. Unpaired Student’s t-test, n = 13.

### TREM2 deficiency differentially modulates glial reactivity in Tg-FDD mouse model

We previously established that in the Tg-FDD model of CAA, an early hallmark of vascular amyloid deposition is the emergence of pronounced astrogliosis occurring in the absence of microglial activation (18). To determine whether endogenous TREM2 reduction differentially affects glial reactivity, we examined the cortex and cerebellum of Tg-FDD and Tg-FDD/TREM2KO mice, quantifying IBA1-positive microglia (Fig. 3A) and GFAP-positive astrocytes (Fig. 3C) within a 120 µm radius around the vasculature. Tg-FDD/TREM2KO mice exhibited a reduced microglial response in the cortex (Fig. 3B), when male and female were analyzed together, whereas microglial activation in the cerebellum remained unchanged. In contrast, astrogliosis was not significantly altered in the cortex, but Tg-FDD/TREM2KO mice exhibited marked astrogliosis in the cerebellum, with elevated GFAP immunoreactivity compared to Tg-FDD male and female mice (Fig. 3D), which is consistent with the increased vascular amyloid accumulation observed in this region. Together, these findings indicate that distinct brain regions exhibit differential glial signatures in response to TREM2 depletion.

**Figure 3:**
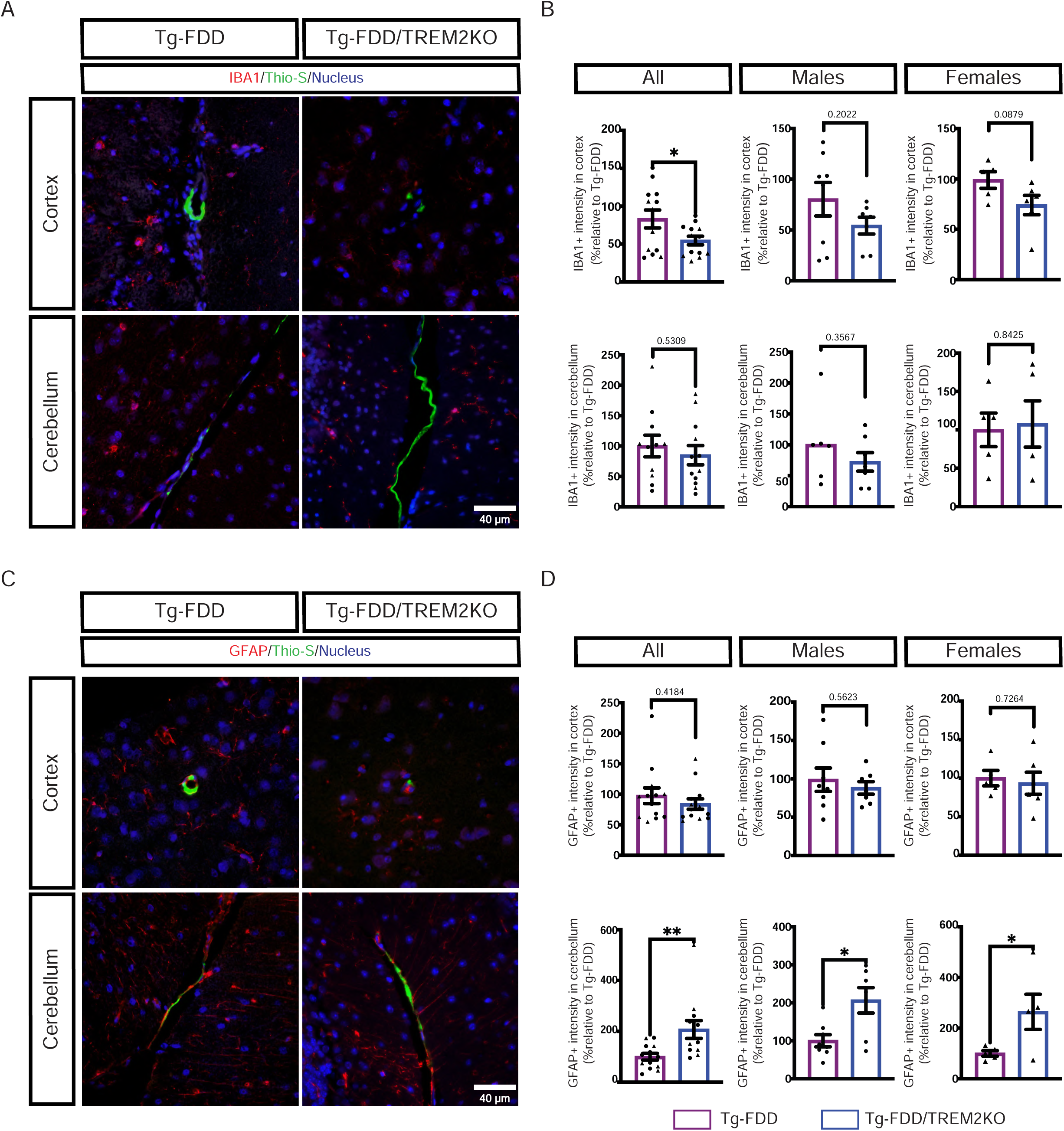
TREM2 depletion enhances astrogliosis in the cerebellum of Tg-FDD mice. Double immunofluorescent staining showing A) amyloid (Thio-S, green) and microglia (IBA1+, red), and C) amyloid (Thio-S, green) and astrocytes (GFAP+, red). Scale bar 40 μm. B) Quantification of IBA1+ intensity and D) Quantification of GFAP+ in the cortex and cerebellum in Tg-FDD and Tg-FDD/TREM2KO mice. Results are shown as the mean ± SEM. Unpaired Student’s t-test, n = 13.

### TREM2 depletion increases tau oligomers in the cerebellum of Tg-FDD mice

Our previous work showed that ADan amyloid induces tau phosphorylation and misfolding, leading to tau-dependent neurotoxicity (28). To assess the spatial relationship between vascular ADan deposits and tau pathology, we performed double immunofluorescent staining int he cortex and cerebellum using the anti-ADan and TOMA1 antibodies (Fig. 4A), which recognize tau oligomers (29,30). In the cortex, double staining showed a reduction of tau oligomers in close proximity to ADan-rich vessels in Tg-FDD/TREM2KO compared with Tg-FDD only in female mice (Fig. 4B). In contrast, in the cerebellum, tau oligomers were increased in Tg-FDD/TREM2KO male and female mice (Fig. 4B). Together, these data indicate that TREM2 leads to distinct perivascular tau pathology in the cortex and cerebellum of Tg-FDD mice, potentially resulting from differential vascular amyloid accumulation.

**Figure 4:**
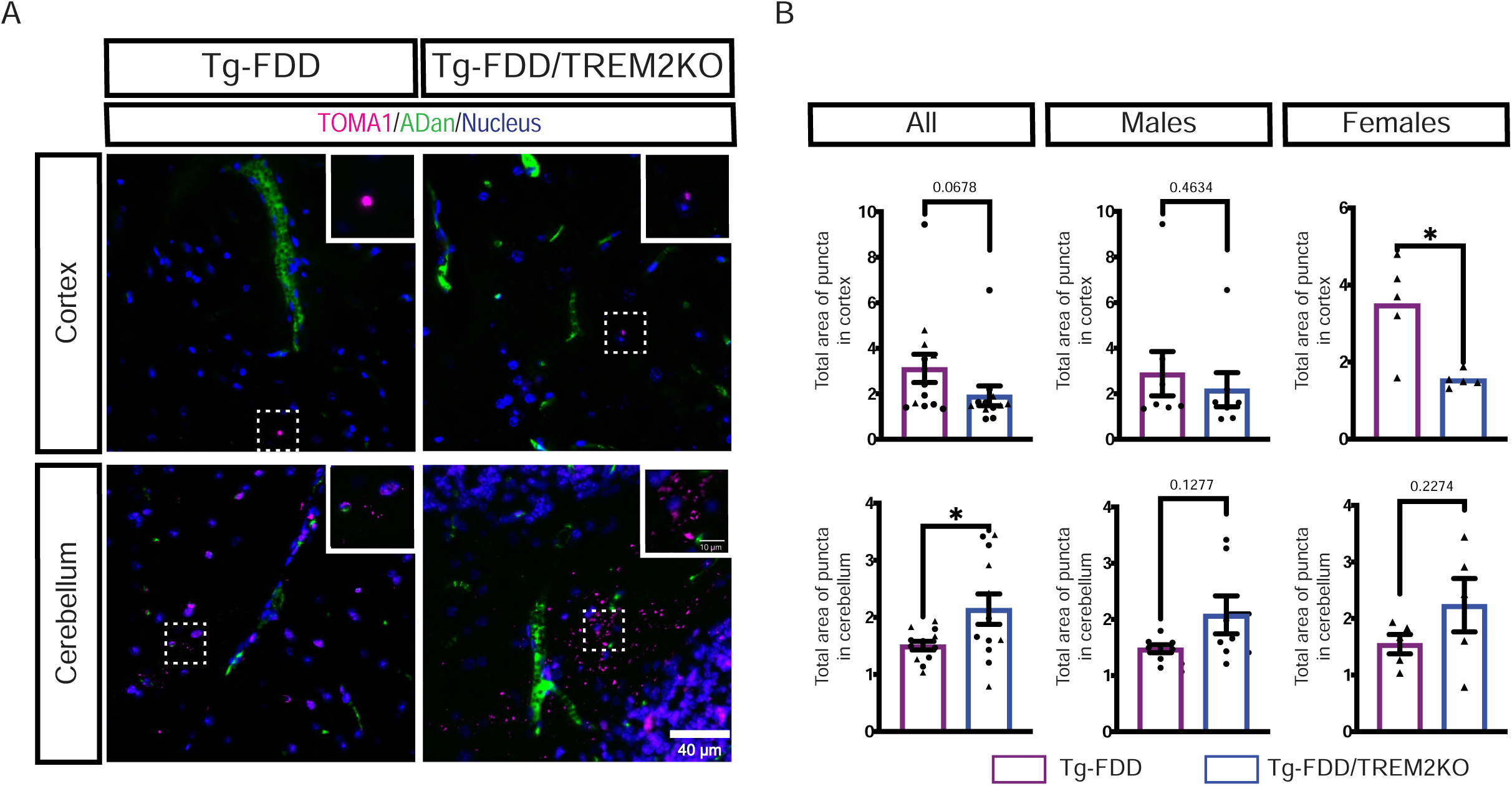
TREM2 deficiency increases tau oligomers in the cerebellum of Tg-FDD mice. Double immunofluorescent staining showing A) amyloid (Adan, green) and tau oligomers (TOMA1, pink). Scale bar 40 μm. B) Quantification of total area of puncta in the cortex and cerebellum in Tg-FDD and Tg-FDD/TREM2KO mice. Results are shown as the mean ± SEM. Unpaired Student’s t-test, n = 13.

### Neuroinflammation pathways exhibit opposite-direction expression in cortex and cerebellum in TREM2-deficient mice within the CAA context

Given the differential outcomes of TREM2 knockout in the cortex and cerebellum in our CAA model, we next investigated whether region-specific transcriptional responses are induced. To do so, we used the Neuroinflammation Panel (NanoString analyses), which quantifies the expression of 770 genes across 23 neuroinflammatory pathways. We evaluated the fold change of every differentially expressed gene (DEG) in all the conditions for both the cortex and cerebellum (Supplementary file 1). The global analysis of DEGs in both brain regions revealed the following. In the cortex (Fig. 5A), green-labeled genes (unique DEGs in the Tg-FDD vs. WT comparison) showed upregulation of epigenetic regulators such as *Dnmt1*, whereas several genes related to immune response were downregulated relative to WT. In contrast, pink-labeled genes (unique DEGs in the Tg-FDD/TREM2KO vs. Tg-FDD comparison) exhibited increased expression of genes associated with immune signaling (*Vav1*), and oligodendrocyte function (*Bcas1*), while key cytokine signaling genes (*Csf3r, Cx3cl1, Il1r1*) were reduced. Genes shared between comparisons (blue-labeled) included transcripts to microglial activity (*Gpr3*4), astrocytic function (*S100b, Nwd1*), extracellular matrix remodeling (*Cldn5, Pecam1, Esam*), and neuronal signaling (*Grm2, Nptx1*).

**Figure 5:**
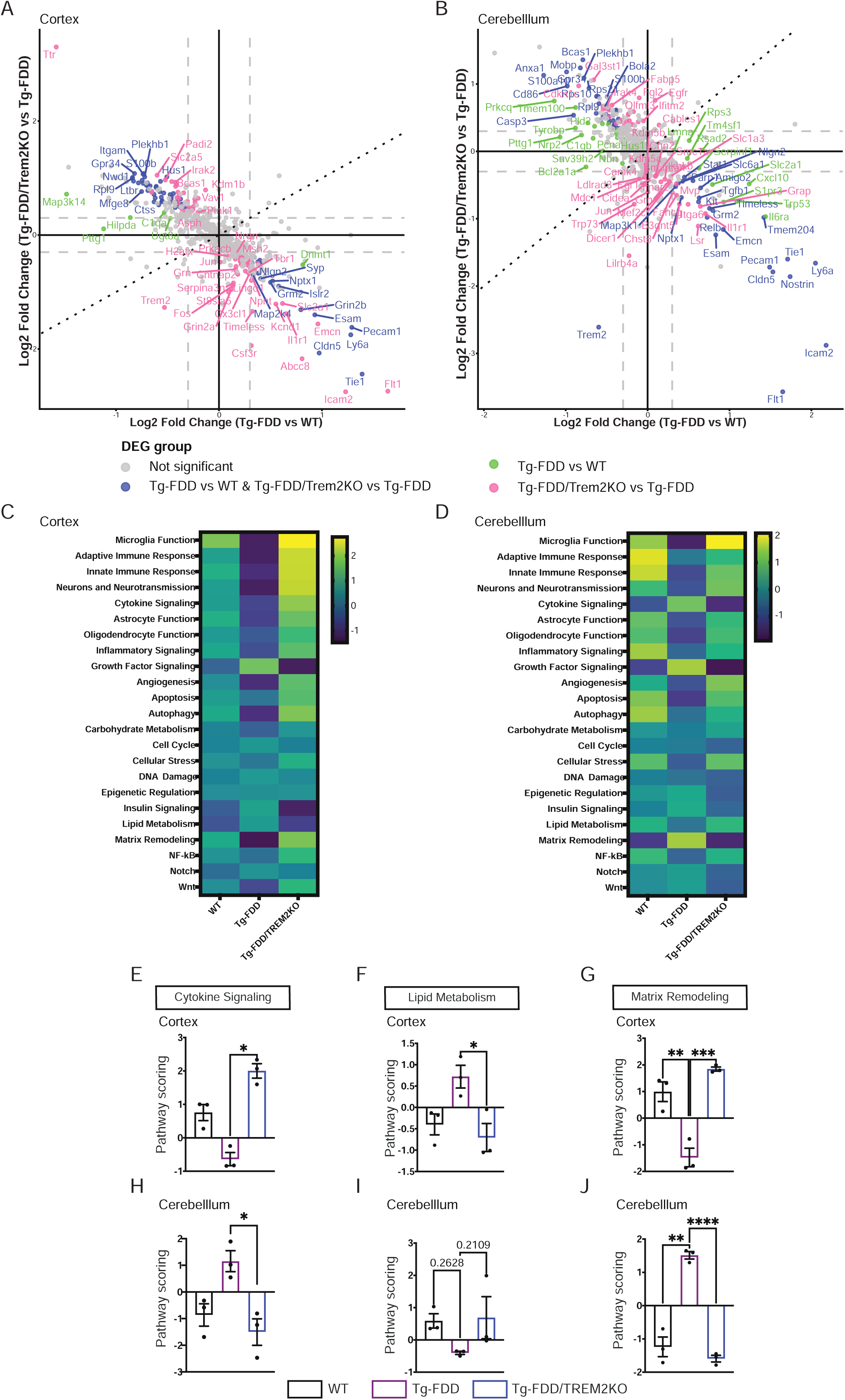
Neuroinflammation pathways exhibit opposite-direction expression in cortex and cerebellum. Total mRNA was isolated from the cortex and cerebellum of wild-type (WT), Tg-FDD, Tg-FDD/TREM2KO mice to perform the NanoString neuroinflammation panel. A, B) “Four-way” plot comparing differential expression in the cortex and cerebellum. Each point represents one gene. The x-axis shows log2 fold changes (Tg-FDD vs WT), and the y-axis shows log2 fold (Tg-FDD vs Tg-FDD/TREM2KO). Genes that are differentially expressed (DEG) with p value of .05 (–log10 (*p*-value) = 1.3) and log2 fold change > 1.2 (log2 (FC) = ± 0.3) in the x-axis comparison are shown in green, the DEGs from the y-axis comparison are shown in pink, and the DEGs in both comparisons are shown in blue. Gray dots represent non-significant genes. C, D) Heatmap showing differential expression of neuroinflammation panel genes in the cortex and cerebellum of WT, Tg-FDD, Tg-FDD/TREM2KO mice. E-J) Pathways score comparisons from neuroinflammation panel. Cytokine signaling, lipid metabolism and matrix remodeling pathways exhibit opposite-direction expression profiles across WT, Tg-FDD, Tg-FDD/TREM2KO mice. Data presented are mean ± SEM. Statistical significance was determined by one-way ANOVA, followed by Tukey’s multiple comparisons. n = 3 per group.

In the cerebellum (Fig. 5B), green-labeled genes in Tg-FDD mice were enriched for immune response pathways (*Tyrobp, Prkcq*), and complement components (*C1qb*), while several microglial-associated genes (*Serpinf1, Slc2a1, Rps3*) showed lower expression. Pink-labeled genes (unique DEGs in Tg-FDD/TREM2KO vs. Tg-FDD) displayed increased expression of matrix remodeling genes (*Olfml3*) and, microglial function genes (*Fabp5*), whereas epigenetic regulators (*Kdm5b*), lipid metabolism genes (*Lsr*), neuronal genes (*Slc1a3*), and cytokine signaling genes (*Il1r1*) were reduced. Blue-labeled genes shared between comparisons were characterized as microglial-related transcripts (*Cd86, Rpl19, Bola2, Rps10, Rps2*), extracellular matrix (*Cldn5, Pecam1, Esam*) and immune response genes (*Icam2, Kit, Stat1*). These results indicate that TREM2 deletion reshapes transcriptional programs in a region-dependent manner, with the cortex showing predominant modulation of cytokine, glial and immune signaling, whereas the cerebellum displays a stronger signature related to matrix remodeling and glial response.

To further explore these effects, we analyzed pathway annotation scores, which integrate gene-level information into a single metric. Distinct expression patterns were observed among the three genotypes in both the cortex and cerebellum, revealing region-specific differences in neuroinflammation-related pathways between WT, Tg-FDD, and Tg-FDD/TREM2KO mice. Notably, three pathways, cytokine signaling, lipid metabolism, and matrix remodeling, showed opposite expression trends between the cortex and cerebellum (Fig. 5C and D), reinforcing the idea that TREM2 deficiency exerts divergent effects on transcriptional programs depending on the brain region.

In the cortex, TREM2 depletion in Tg-FDD mice increased both cytokine signaling and matrix remodeling pathway scores (Fig. 5E and G), whereas Tg-FDD mice showed significantly reduced matrix remodeling scores compared to WT (Fig. 5G). In contrast, lipid metabolism pathway scores were decreased in Tg-FDD/TREM2KO mice relative to Tg-FDD animals (Fig. 5F). In the cerebellum, cytokine signaling and matrix remodeling pathway scores were reduced in Tg-FDD/TREM2KO compared to Tg-FDD mice (Fig. 5H and J), while Tg-FDD mice exhibited significantly increased matrix remodeling relative to WT. Conversely, lipid metabolism pathway scores tended to decrease in Tg-FDD mice compared to WT, with a trend toward increased scores in Tg-FDD/TREM2KO mice (Fig. 5I). Overall, TREM2 deficiency is associated with region-specific changes in pathway activity, with opposing patterns observed between the cortex and cerebellum.

### Expression profiles of three neuroinflammation pathways differ between cortex and cerebellum within the CAA context

We next analyzed DEGs related to cytokine signaling, lipid metabolism, and matrix remodeling in the cortex and cerebellum of WT, Tg-FDD, and Tg-FDD/TREM2KO mice. In the context of cytokine signaling pathway (Fig. 6A and B), in the cortex, Tg-FDD mice (green-labeled) showed reduced *Map3k14*, whereas TREM2 depletion in Tg-FDD mice (pink-labeled) further elevated *Csf1r*, *Tnfsf12*, *Irak2*, *Tgfbr1*, *Traf3,* and *Vav1*. Shared DEGs (blue-labeled) included *Cx3cr1*, *Akt2*, and *Ltbr* were increased in Tg-FDD/TREM2KO and reduced Tg-FDD (Fig. 6A). In the cerebellum, Tg-FDD mice showed downregulation of *Pik3r1* and upregulation of *Cxcl10* and *Il6ra* compared to WT (green dots). In Tg-FDD/TREM2KO mice, *Irak4* and *Traf3* were specifically increased relative to Tg-FDD (pink dots), while *Sumo1* was elevated in Tg-FDD/TREM2KO but reduced in Tg-FDD. Conversely, *Kit* and *Stat1* showed increased expression in Tg-FDD and decreased expression in Tg-FDD/TREM2KO (blue dots) (Fig. 6B). For lipid metabolism–related genes, *Apoe* was decreased in Tg-FDD compared to WT and increased with TREM2 depletion in Tg-FDD mice, in both the cortex and cerebellum (Fig. 6C and D).

**Figure 6:**
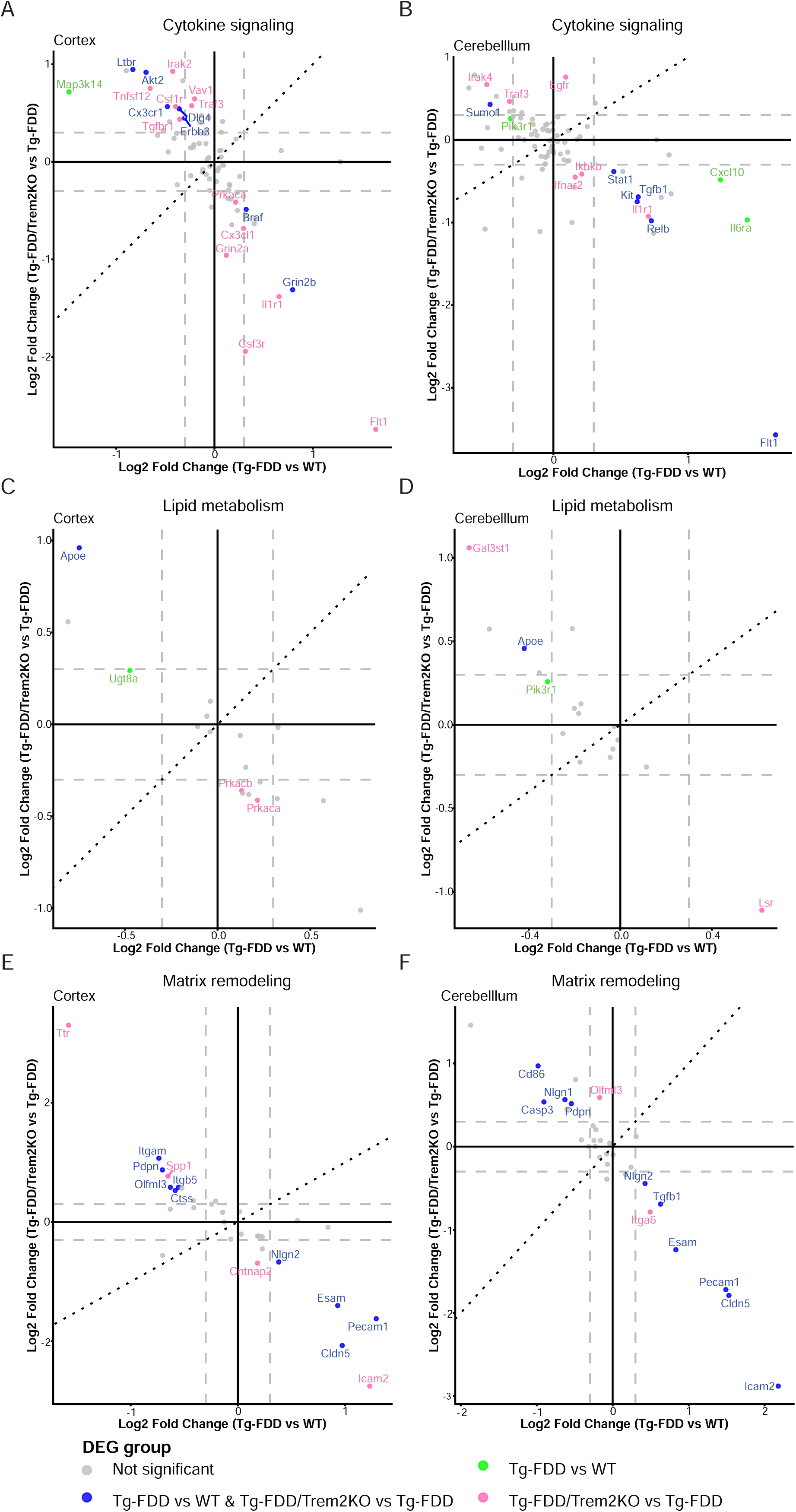
Expression profiles of three neuroinflammation pathways show differences between cortex and cerebellum. Total mRNA was isolated from the cortex and cerebellum of wild-type (WT), Tg-FDD, Tg-FDD/TREM2KO mice and analyzed using the NanoString neuroinflammation panel. “Four-way” plot comparing differential expression in the cortex and cerebellum: A, B) cytokine signaling; C, D) lipid metabolism; E, F) matrix remodeling. Each point represents one gene. The x-axis shows log2 fold changes (Tg-FDD vs WT), and the y-axis shows log2 fold (Tg-FDD vs Tg-FDD/TREM2KO). Genes that are differentially expressed (DEG) with p value of .05 (–log10 (*p*-value) = 1.3) and log2 fold change > 1.2 (log2 (FC) = ± 0.3) in the x-axis comparison are shown in green, the DEGs from the y-axis comparison are shown in pink, and the DEGs in both comparisons are shown in blue. Gray dots represent non-significant genes. n = 3 per group.

Regarding extracellular matrix remodeling, in the cortex, Tg-FDD/TREM2KO mice (pink-labeled) selectively upregulated *Spp1* and *Ttr* and downregulated *Icam2* compared to Tg-FDD. Genes upregulated (blue-labeled) in Tg-FDD/TREM2KO, including *Itgam*, *Pdpn*, *Olfml3*, *Itgb5*, and *Ctss,* showed the opposite pattern in Tg-FDD. Conversely, *Cldn5, Pecam1, Nlgn2, and Esam* were downregulated in Tg-FDD/TREM2KO and upregulated in Tg-FDD (Fig. 6E). In the cerebellum, *Olfml3* was increased in Tg-FDD/TREM2KO, whereas *Itga6* was decreased (pink dots). Shared genes across both comparisons (blue dots), including *Cd86*, *Nlgn1*, *Casp3*, and *Pdpn,* were upregulated in Tg-FDD/TREM2KO but downregulated in Tg-FDD. In contrast, *Esam*, *Pecam1*, and *Cldn5* were downregulated in Tg-FDD/TREM2KO and upregulated in Tg-FDD (Fig. 6F). These findings indicate that TREM2 loss reshapes transcriptional programs in a region- dependent manner, with distinct effects on cytokine signaling, lipid metabolism, and matrix remodeling, in the cortex versus the cerebellum.

## DISCUSSION

Alterations in TREM2 function have been linked to increased amyloid deposition and dysregulated inflammation, supporting its therapeutic relevance (31). Here, we provide the first *in vivo* evidence that TREM2 deletion exerts distinct, region-specific effects in a mouse model of CAA. Loss of endogenous TREM2 differentially modulated vascular amyloid accumulation, glial responses, and tau pathology in the cortex and cerebellum, accompanied by distinct inflammatory profiles. These findings highlight the importance of developing therapeutic strategies adapted to the regional complexity of CAA/AD.

In our study, TREM2 ablation in Tg-FDD mice produced region-specific effects on amyloid deposition, increasing vascular amyloid in the cerebellum while reducing cortical burden. Similar heterogeneity has been reported in other AD and CAA models after TREM2 deficiency: in the APPPS1 model, which shows robust parenchymal Aβ plaque deposition, Aβ burden varies across regions (32), and in Tg-SwDI mice, which predominantly develop CAA, exhibit increases parenchymal amyloid alongside reducing CAA (33). Overall, these findings indicate that the impact of TREM2 on amyloid pathology, depend on both brain region and the underlying amyloid context.

Previous studies have shown that the brain responds to pathology in a region-specific manner, with distinct inflammatory responses across areas such as the cortex, cerebellum, and hippocampus (34–36). Consistent with this, our NanoString analysis revealed distinct inflammatory profiles between brain regions in Tg-FDD/TREM2KO mice, indicating that TREM2 deficiency drives region-specific transcriptional programs. Notably, cytokine signaling, lipid metabolism, and matrix remodeling pathways showed opposite patterns across cortex and cerebellum, reinforcing this regional specificity.

Cytokine signaling pathway scoring showed a global increase in the cortex of Tg-FDD/TREM2KO mice, accompanied by decreased expression of *Cx3cl1* and *Prkaca*, two key regulators of microglial homeostasis (37,38). CX3CL1–CX3CR1 signaling supports microglial responsiveness and a pro-inflammatory, phagocytic phenotype, and its disruption impairs microglial function (37,39). Similarly, PRKACA-mediated cAMP–PKA signaling suppresses pro-inflammatory cytokine production and promotes IL-10 expression in microglia (38), suggesting reduced regulatory control of cytokine responses. In contrast, the cerebellum, where amyloid load and astrogliosis were higher, showed an opposite cytokine signaling pattern, alongside increased expression of *Pik3r1*, a component of the PI3K–AKT pathway implicated in Aβ/tau pathology and astrocyte reactivity (40,41). Notably, TREM2 has been proposed as a key regulator of microglial–astrocyte communication required for proper glial crosstalk and astrocytic responses (42), and its loss may alter intercellular signaling between microglia and astrocytes. These findings suggest that TREM2 regulates region-specific inflammatory states through distinct signaling pathways in cortex and cerebellum.

Matrix remodeling pathways also showed opposite scoring between regions. Interestingly, matrix composition varies across brain regions in both AD and CAA, reflecting local vascular stress and amyloid burden (43). In the cortex, this pathway scoring was reduced in Tg-FDD mice and restored upon TREM2 depletion. Specifically, Tg-FDD/TREM2KO mice showed increased expression of genes associated with microglial recruitment, phagocytic activity, inflammatory signaling, and matrix interaction, including *Itgam*, *Itgb5*, and *Ctss* (44–46), together with reduced *Pecam1*, a marker of vascular integrity (47). In contrast, the cerebellum exhibited increased *CD86* expression, consistent with a more reactive astrocytic state (74). Together, these region-specific differences suggest that TREM2-dependent regulation of microglia–astrocyte communication is shaped by the local brain microenvironment, leading to distinct glial states and matrix remodeling responses in cortex and cerebellum, recapitulating the effects observed in cytokine signaling.

Lipid metabolism pathways were also affected in a region-specific manner. Vascular amyloid deposits strongly interact with lipids, promoting Aβ aggregation and retention in cerebral vessels (48), and regional differences in lipid composition may further influence these processes (49). *APOE*, a key regulator of brain lipid transport, modulates vascular Aβ clearance and interacts closely with TREM2 signaling (50–52), which has been shown to directly regulate its expression (52). Consistently, TREM2 depletion increased *Apoe* levels in both regions, supporting activation of a direct TREM2–APOE regulatory program. This pathway has been reported to define microglial transcriptional states in neurodegeneration and to shape downstream astrocytic responses (42,51), providing a mechanistic framework for the lipid-related alterations observed. These results suggest that altered lipid metabolism may contribute to the region-specific effects of TREM2 loss.

Finally, we observed region-dependent effects of TREM2 on tau pathology. In Tg-FDD mice, TREM2 deficiency had region-specific effects on tau, perivascular oligomers decreased in the cortex but increased in the cerebellum, consistent with regional amyloid deposits. Interestingly, our laboratory demonstrated that tau mediates neurotoxicity in CAA (28). TREM2 loss can exacerbate tau accumulation and propagation, particularly near Aβ plaques, while in tauopathy models it generally promotes tau phosphorylation and aggregation (7,13,53–55). Recent studies further support region-specific effects of TREM2, with differential tau accumulation, phosphorylation, and aggregation observed across brain regions in various models following TREM2 depletion (13,56). Together, these findings support a context and region dependent role of TREM2 in modulating tau pathology.

Overall, our study provides *in vivo* evidence that TREM2 deficiency exerts region-specific effects in CAA. These findings highlight the role of TREM2 in coordinating neuroinflammatory responses to vascular amyloid pathology and suggest that therapeutic strategies targeting TREM2 in Alzheimer’s Disease and Related Dementias (ADRD) should consider its effect on vascular pathology as well as brain-region specificity.

## AUTHOR CONTRIBUTIONS

C.L-R. designed the study and C.M and C.L-R wrote the manuscript. C.M. designed and performed the experiments. R.V. provided Tg-FDD mice. N.J-G and J.M-P assisted in animal procedures. A.A. assisted with figure preparation and data analysis. N.J-G. and A.A. provided scientific guidance and data analysis. C.M, N.J-G and C.L-R critically revised the manuscript and interpreted the data. All the authors have read and approved the final manuscript.

## ACKNOWLEDGMENTS

This research was supported by the National Institute of Health and the National Institute for Neurological Disorders and Stroke (NIH/NINDS) 1R01NS119280, the National Institute of Health and the National Institute of Aging (NIH/NIA) 1RF1AG059639 grants, and ALZDISCOVERY-1049108 from the Alzheimer’s Association grant; the Cure Alzheimer’s Fund; and the Rainwater Charitable Foundation to Cristian Lasagna-Reeves. NanoString data were generated in part through the use of the Advanced Technology Genomics Core (ATGC), which receives partial support from the National Cancer Institute under grant CA016672 to MD Anderson Cancer Center.

## CONSENT STATEMENT

No human subject materials were utilized in this study.

## Supplementary

**Supplementary Table 1: RNA quality parameters for NanoString analysis**. Total mRNA was isolated from the cortex and cerebellum of 22-month-old male WT, Tg-FDD, and Tg-FDD/TREM2KO mice. RNA quality was assessed by DV200 (%) and RNA Integrity Number (RIN) values.

**Supplementary file 1:** List of differentially expressed genes (DEGs) and normalized counts in the cortex and cerebellum of Tg-FDD, WT, and Tg-FDD/TREM2KO mice.

