## Supplementary material for "TREM2 deficiency causes region-specific brain effects in a mouse model of cerebral amyloid angiopathy": Supp. Table 1

| Samples | Concentration ng/µl | DV 200% | RIN |
|---|---|---|---|
| Tg-FDD 1 cortex | 1680 | 95.73 | 8.4 |
| Tg-FDD 1 cerebellum | 368 | 93.95 | 8.9 |
| Tg-FDD 2 cortex | 1340 | 94.21 | 8.7 |
| Tg-FDD 2 cerebellum | 660 | 93.85 | 8.8 |
| Tg-FDD 3 cortex | 1760 | 95.01 | 8.5 |
| Tg-FDD 3 cerebellum | 480 | 94.22 | 8.9 |
| WT 1 cortex | 1300 | 93.11 | 8.8 |
| WT 1 cerebellum | 438 | 91.86 | 8.8 |
| WT 2 cortex | 880 | 95.19 | 8.7 |
| WT 2 cerebellum | 310 | 92.15 | 8.9 |
| WT 3 cortex | 1000 | 93.44 | 8.8 |
| WT 3 cerebellum | 412 | 91.6 | 8.8 |
| Tg-FDD/TREM2KO 1 cortex | 1260 | 93.36 | 8.8 |
| Tg-FDD/TREM2KO 1 cerebellum | 420 | 92.82 | 8.9 |
| Tg-FDD/TREM2KO 2 cortex | 1280 | 94.42 | 8.7 |
| Tg-FDD/TREM2KO 2 cerebellum | 318 | 89.63 | 8.9 |
| Tg-FDD/TREM2KO 3 cortex | 1200 | 94.97 | 8.7 |
| Tg-FDD/TREM2KO 3 cerebellum | 426 | 93.23 | 8.7 |
